# 3D Cell-Matrix Mechanical Interaction Models for Cancer Invasion and Drug Evaluation

**DOI:** 10.64898/2026.02.02.703199

**Authors:** Xinyu Jin, Jing Jiao, Cheng Qian, Bingqi Ning, Zimeng Zhang, Haoxiang Zhang, Liang Qiu, Ran Zhang, Susana Rocha, Hailong Wang, Chao Fang, Chengfen Xing, Hongbo Yuan

## Abstract

Cancer cells breach the extracellular matrix (ECM) using both protease-driven degradation and force-driven physical remodeling, yet most anti-metastatic drug screens still rely on biochemical assays that overlook cell–matrix mechanical reciprocity. Here, we present a fully synthetic 3D invasion platform based on cellular force-responsive polyisocyanide (PIC) hydrogels that isolates biophysical invasion mechanisms. Cell-generated forces align and densify the PIC fibrous network, reproducing hallmark matrix remodeling seen in the tumor microenvironment. A constitutive model, parameterized by the critical stress for strain stiffening effect, links matrix nonlinear elasticity to pericellular stiffening, long-range mechanotransmission, and intercellular coupling. Using this system, we show that breast cancer cells invade by pulling and pushing the network even when matrix metalloproteinases are inhibited, revealing a physical bypass of protease blockade. Accordingly, broad-spectrum metalloproteinase inhibitors that suppress invasion in Matrigel fail to inhibit invasion here, exposing a limitation of current drug-evaluation pipelines. In co-culture, cancer-associated fibroblasts markedly accelerate invasion by generating aligned fiber tracks through higher contractility, implicating CAF-driven mechanical remodeling as a key route for breaching barriers during metastasis. The platform is thermoresponsive, compatible with standard Transwell formats, enables direct imaging of fiber architecture and invasion fronts, and decouples biophysical from biochemical cues for mechanism-aware, animal-free assessment of anti-metastatic therapies.

## Main

To metastasize, cancer cells must surpass the physical hurdles imposed by surrounding extracellular matrix (ECM). Increasing evidence shows that both biochemical and biophysical modes of matrix remodeling are critical in this process. Tumor cells degrade the ECM biochemically through secretion of matrix metalloproteinases (MMPs), which allows them to breach the basement membrane. In parallel, tumor cells and cancer-associated fibroblasts (CAFs) remain physically coupled to the matrix. CAFs, in particular, exert contractile forces on the basement membrane, leading to fiber alignment and enlargement of pre-existing pores. These changes facilitate tumor cell invasion through an MMP-independent mechanism^1-4^.

Given the significant roles of matrix remodeling in both pro-tumorigenesis and anti-tumorigenesis, targeting matrix remodeling has been considered as a powerful approach in recent years^4,5^. For example, inhibition of enzymatic crosslinking by lysyl oxidasesor transglutaminases reduces desmoplasia, softens the stromal barrier, and improves therapeutic response in preclinical models of pancreatic, colorectal, and breast cancers^6-9^. MMPs also play central roles in invasion, angiogenesis, and metastasis, suggesting that broad inhibition of these enzymes should be effective^10^. Accordingly, MMP inhibitors have been pursued as anticancer drugs, but clinical outcomes have been disappointing. The broad-spectrum MMPs inhibitor, Marimastat failed to improve survival either as a monotherapy or in combination with gemcitabine in unresectable pancreatic cancer, and it showed only modest benefit in non-small-cell lung cancer when combined with carboplatin and paclitaxel^11-13^. Similarly, Tanomastat, which targets MMP-3, MMP-9, and MMP-13, demonstrated poor efficacy in phase III trials in pancreatic cancer patients treated with gemcitabine^14^. These results underscore the complexity of matrix remodeling in metastasis and highlight the need for models that account for both biochemical and mechanical contributions to tumor progression.

The Transwell (Boyden chamber) invasion assay is one of the most widely used in vitro methods for evaluating cell migration and invasion across extracellular matrix barriers, particularly in cancer and immune cell research^15,16^. In its standard configuration, a porous membrane is coated with a thin layer of Matrigel, cells are seeded on top in serum-free medium, and a chemoattractant in the lower chamber establishes a gradient. Cells that traverse the matrix barrier and reach the underside of the membrane are then quantified, providing a readout of migratory or invasive capacity in response to experimental cues. While simple and convenient, this assay has important limitations for mechanistic studies. Matrigel is derived from mouse tumors and contains a highly complex and poorly defined mixture of proteins and growth factors^17^. Moreover, critical extracellular parameters such as stiffness, fiber architecture, and pore size cannot be systematically tuned^18^. As a result, the assay provides limited insight into the underlying cell–matrix interactions and has reduced predictive power for therapeutic testing.

Engineered hydrogels provide versatile platforms to study how the extracellular matrix regulates cell behavior in vivo and to create three-dimensional (3D) systems that intentionally guide migration for biomedical applications^18-20^. Synthetic hydrogels such as poly(ethylene glycol) and dextran have been functionalized with protease-cleavable crosslinks, enabling cells to locally degrade the network and migrate through it^21-23^. In native tissues, however, invasion is not limited to enzymatic degradation. Cells can also reorganize the surrounding matrix through contractile forces, exploiting the plasticity of fibrous ECM to generate persistent deformations that form migration channels^2,24-26^. The fibrous and anisotropic architecture of ECM plays a central role in this process^27,28^. For example, cells align and densify fibers along their migration axis, creating tracks that act as “highways” to direct movement and promote collective invasion^29,30^. Computational models further suggest that the nonlinear mechanics of fibrous matrices facilitate long-range force transmission and cell–cell communication, providing a rapid and efficient signaling mechanism in 3D that complements biochemical pathways^31-33^.

Recently, we reported a highly biomimetic hydrogel based on polyisocyanide (PIC) polymers, which combines fibrous architecture with nonlinear mechanical properties^34,35^. PIC hydrogels respond sensitively to cellular forces and replicate essential aspects of ECM–cell reciprocity, including fiber alignment, matrix densification, and long-range force propagation, underscoring their potential as a powerful tool for deciphering cell–matrix mechanoreciprocity^36,37^.

Here, we established a 3D cancer invasion model based on PIC hydrogels to decouple biochemical from biophysical cell–matrix interactions (**Figure 1**). Owing to their fibrous architecture and nonlinear mechanical response, PIC matrices dynamically adapt to cell-generated forces, supporting bidirectional interactions in which cells align and densify fibers to create migration tracks that closely mimic features of the tumor microenvironment. Using this system, we observed that CAFs promote cancer cell invasion through mechanical remodeling of the matrix, even in the absence of MMP activity. Consistently, broad-spectrum MMP inhibitors that appeared effective in Matrigel failed to suppress invasion in PIC, indicating that physical remodeling can bypass enzymatic inhibition. To mechanistically interpret these behaviors, we developed a constitutive model of the PIC network with tunable mechanical parameters and applied it to probe how matrix mechanics influences cell–cell communication. The simulations reproduced experimental observations and revealed how nonlinear fibrous mechanics facilitate long-range mechanotransmission that supports coordinated invasion and metastatic spread. The platform is thermoresponsive, animal-free, and compatible with standard workflows, providing protocols and analysis that decouple matrix chemistry from mechanics. Together, these findings underscore the importance of evaluating both biochemical and biophysical pathways of cell–matrix interaction in order to better understand tumor invasion and to improve the predictive value of preclinical drug testing.

**Figure 1.**
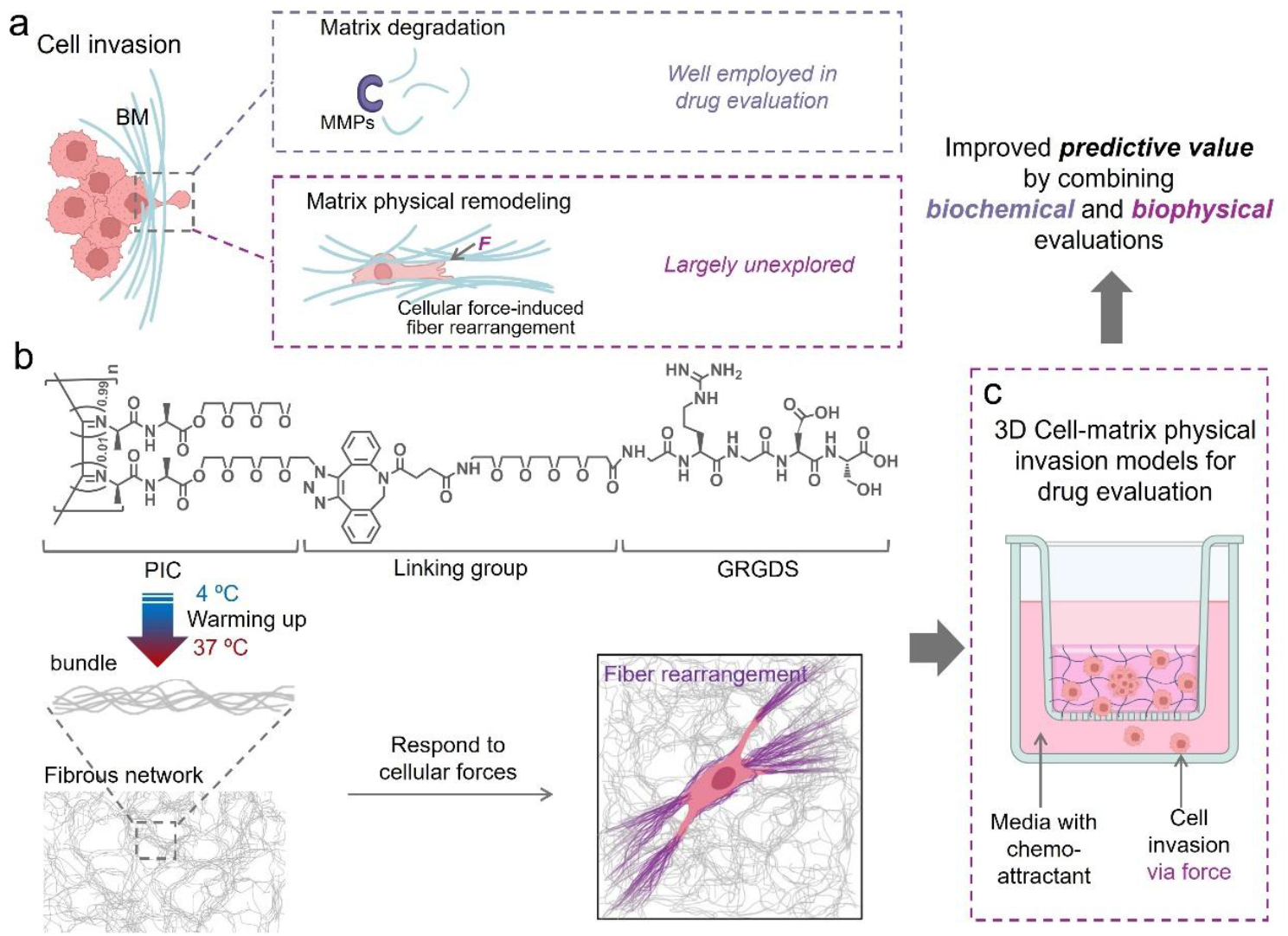
PIC-based 3D invasion model captures physical matrix remodeling for drug evaluation. (a) Schematic of two major cancer cell invasion strategies: biochemical degradation of the basement membrane (BM) by MMPs, which is widely studied and used in drug evaluation, and physical matrix remodeling via cellular force– induced fiber rearrangement, which remains largely unexplored. (b) Chemical structure of PIC functionalized with the integrin-binding peptide GRGDS. Upon warming, PIC assembles into fibrous bundles that form a 3D network (Lower left panel). The fibrous matrix responds to cellular forces, enabling fiber alignment and densification that mimic natural ECM remodeling (Right lower panel). (c) PIC hydrogel-based Transwell assay for drug evaluation. A thin PIC/cell layer is placed on the insert with chemoattractant in the lower chamber. Invasion occurs via force-mediated remodeling, allowing assessment of biophysical contributions to invasion and drug response.

## Results

### Tunable PIC mechanics regulate cancer cell and CAF behaviors

To generate matrices that respond across different ranges of cellular force, we prepared two classes of PIC hydrogels with distinct mechanical properties. Both the linear and nonlinear mechanics of PIC can be tuned by polymer concentration *c* and, at fixed *c*, by polymer length^38,39^. PIC is thermoresponsive and reversible, undergoing a rapid sol– gel transition when the temperature exceeds the gelation point, and reverting to a sol upon cooling. This property enables straightforward cell encapsulation, retrieval, and integration with downstream assays. As shown in **Figure 2**a, the storage modulus *G*′ increases with temperature, and at *c* = 1.0 mg mL^−1^, longer polymers produce stiffer gels with a slightly reduced gelation temperature. Nonlinear behavior was probed using a prestress protocol in which a constant stress deforms the gel while a superimposed oscillatory stress measures the shear response, yielding the differential modulus *K*′= *∂σ/∂γ*, which is the instantaneous rate of change of shear stress (*σ*) with respect to shear strain (*γ*)^40^. At low stress, *K′* corresponds to the plateau modulus *G*_0_, but rises steeply beyond the critical stress *σ*_c_, consistent with the strain-stiffening response of collagen hydrogels (**Figure 2**b). Importantly, *σ*_c_ serves as an indicator of the matrix’s sensitivity to external or cell-generated forces, and can be tuned by polymer length^41^. Longer PIC polymers exhibit higher *σ*_c_, thereby raising the force threshold for matrix recruitment. Because matrix plasticity is a critical determinant of remodeling and invasion, we next performed macroscopic creep–recovery experiments to quantify the plasticity of PIC hydrogels. Under constant load, PIC displayed time-dependent strain, followed by partial recovery after stress release. As shown in **Figure 2**c, short PIC deformed more readily and recovered less compared with long PIC hydrogels, indicating a higher degree of plasticity. This behavior suggests that short polymer networks are more permissive to persistent cell-induced remodeling.

**Figure 2.**
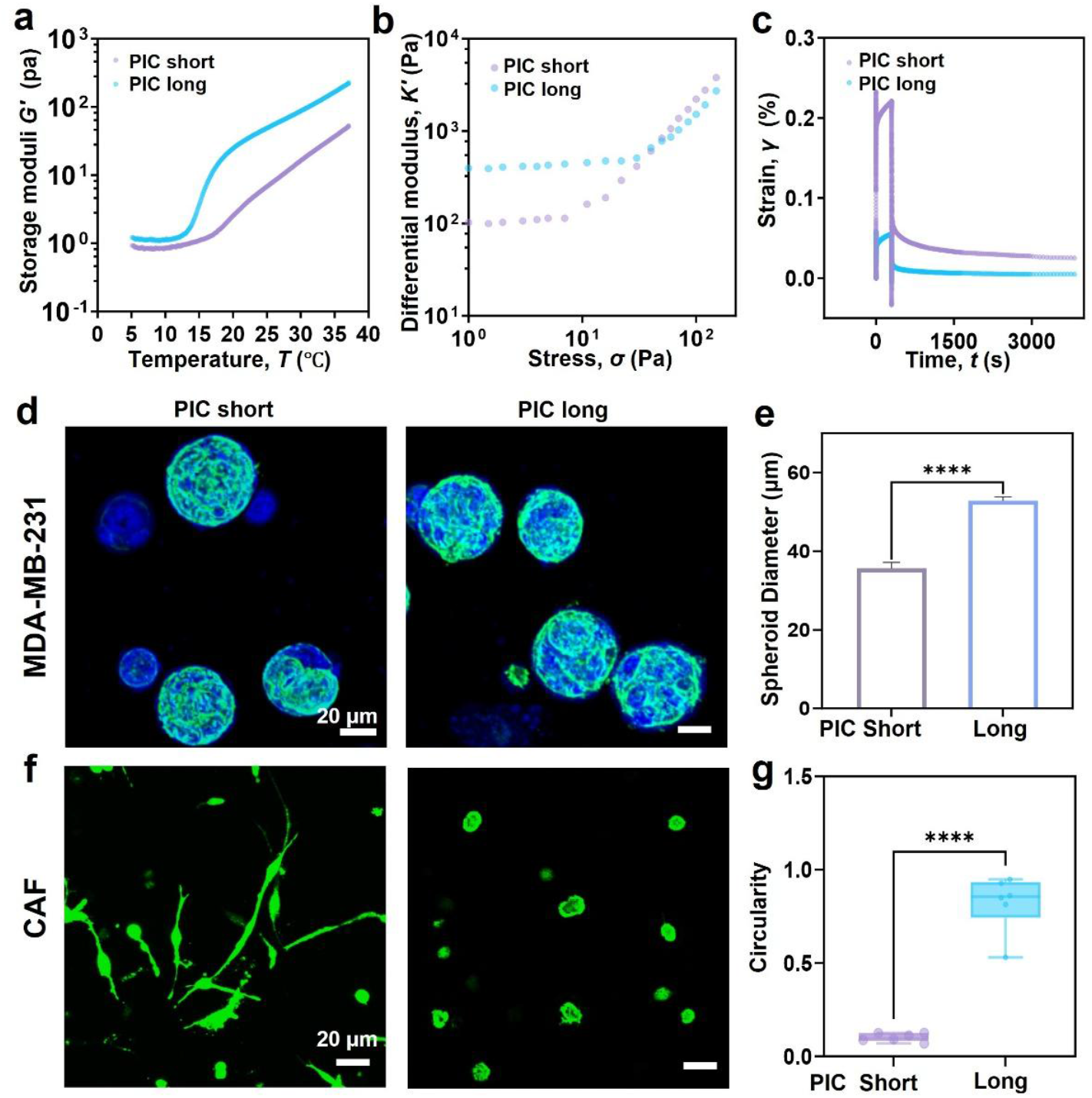
Mechanical characterization of PIC hydrogels and corresponding cell responses. (a) Storage moduli (*G*′) of PIC hydrogels as a function of temperature for different polymer lengths (short and long, see Experimental Section), showing thermoresponsive gelation. (b) Differential modulus (*K*′) versus applied stress, demonstrating a strain-stiffening effect with a higher critical stress (onset of the nonlinear regime) in longer PIC. (c) Creep–recovery tests reveal greater irreversible deformation (plasticity) in short compared with long PIC. PIC hydrogel samples for mechanical characterization were prepared in PBS at 1.0 mg mL^-1^. (d) Confocal images of MDA-MB-231 breast cancer cells forming spheroids in short and long PIC hydrogels with GRGDS functionalization after 7 days. Nuclei: DAPI (blue); F-actin: phalloidin (green). (e) Quantification of spheroid diameters shows significantly larger spheroids in long PIC. (f) Confocal images of GFP-expressing CAFs (green) morphology in short and long PIC hydrogels with GRGDS functionalization after 1 day. (g) Circularity analysis indicates extensive CAF spreading in short PIC, whereas cells remain rounded in long PIC. PIC hydrogels used in panels d–g were functionalized with GRGDS at a polymer concentration of 1.0 mg mL^-1^, corresponding to a GRGDS density of 31.4 μM. Data represent the mean ± SEM (n = 6-10, ****p < 0.0001, One-way ANOVA Tukey’s test). Scale bar = 20 μm.

We employed a highly metastatic breast cancer cell line (MDA-MB-231) together with immortalized CAFs to establish an in vitro model and investigate cell–matrix mechanical interactions during metastasis. To support adhesion, PIC polymers were functionalized with 1% of the integrin-binding peptide GRGDS, corresponding to a peptide density of 31.4 μM at a polymer concentration of 1.0 mg mL^−1^, consistent with earlier reports showing this modification is sufficient for cell attachment and growth^35^. As shown in **Figure 2**d and **Figure S1** in the supporting information, MDA-MB-231 cells proliferated and progressively assembled into multicellular spheroids within RGD-functionalized PIC hydrogels over 7 days. Longer polymer PIC hydrogels, which display higher stiffness and critical stress, promoted more pronounced proliferation and yielded larger spheroids (**Figure 2**e). In contrast, PIC without adhesion ligand supported only small, loosely organized spheroids (**Figure S2** in the supporting information). These observations are consistent with established roles of matrix stiffness in tumor progression and underscore the importance of both biochemical cues and mechanical properties in shaping tumor cell growth.

Within the tumor microenvironment, CAFs are persistently activated fibroblasts that generate strong contractile forces and contribute to tumor progression, metastasis, and therapeutic resistance by secreting soluble factors and remodeling the extracellular matrix^42-44^. While many studies have highlighted the effects of CAF-derived cytokines and growth factors on cancer cells, here we focus on the mechanical remodeling and cell–cell communication mediated by CAFs. As shown in **Figure 2**f and **Figure S3** in the supporting information, CAFs displayed distinct morphologies in both short- and long-polymer RGD-functionalized PIC hydrogels after 1 day of culture. Cell spreading was quantified by measuring circularity (**Figure 2**g). Although the storage moduli of both hydrogel types fall within the soft biomaterial range, differences in the critical stress *σ*_c_ determined how cells perceived their mechanical environment. A lower *σ*_c_ in short PIC increased network sensitivity to cell-generated forces, allowing relatively small contractile inputs to trigger pronounced stiffening. This effect was further enhanced by the high stiffening index, enabling traction forces to rapidly recruit additional fibers into load-bearing. Consequently, in short PIC, CAFs readily engaged the fibers, generated sufficient traction, and underwent spreading and morphological transitions. By contrast, long PIC hydrogels exhibited higher *σ*_c_, so although adhesion was maintained, CAFs were unable to accumulate enough force to overcome the stiffening threshold and therefore remained rounded. Importantly, CAFs still maintained high viability in short PIC after 7 days (**Figure S4** in the supporting information).

### Matrix stiffening facilitates long range intercellular communications

PIC hydrogels recapitulate the two essential features of fibrous ECM, architecture and nonlinear mechanics, and, uniquely, permit modulation of the critical stress for strain-stiffening (*σ*_c_) by adjusting polymer length. To elucidate how variations in polymer length determine the behaviors and morphology of embedded cells, we resorted to theoretical modeling. Actually, continuum models for polymers have progressed from isotropic hyperelastic formulations to fiber-reinforced, tension-only constitutive laws with a strain-stiffening switch^32,33^. Here, we formulated PIC as an isotropic background matrix coupled with a recruited fibrous network, following a constitutive law originally developed for collagen network^32^. The total Helmholtz free-energy density is expressed as

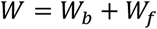

with *W*_*b*_ and *W*_*f*_ being the contribution from isotropic response and aligned fibers, respectively. Specifically, *W*_*b*_ takes a Neo-Hookean form, while *W*_*f*_ is characterized with a critical stretch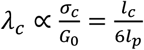 (Supporting Information), where *l*_*c*_ and *l*_*p*_ are the contour and persistence lengths of the polymer, respectively. The principal stresses arising from the fiber alignment vanish below this critical stretch, and exhibit a stiffened response beyond it.

With this model, bulk simple-shear simulations reproduced the experimentally measured nonlinear stress–strain response of PIC hydrogels (**Figure 3**a). Moreover, previous bulk rheology measurements demonstrated that increasing the PIC polymer contour length *l*_*c*_ gradually raises the network’s nonlinear critical stress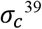, which in turn affects the critical stretch λ_*c*_. By tuning λ_*c*_, our model accurately captured stress– strain curves for PIC hydrogels with different polymer lengths, in agreement with experimental data (**Figure 3**b).

**Figure 3.**
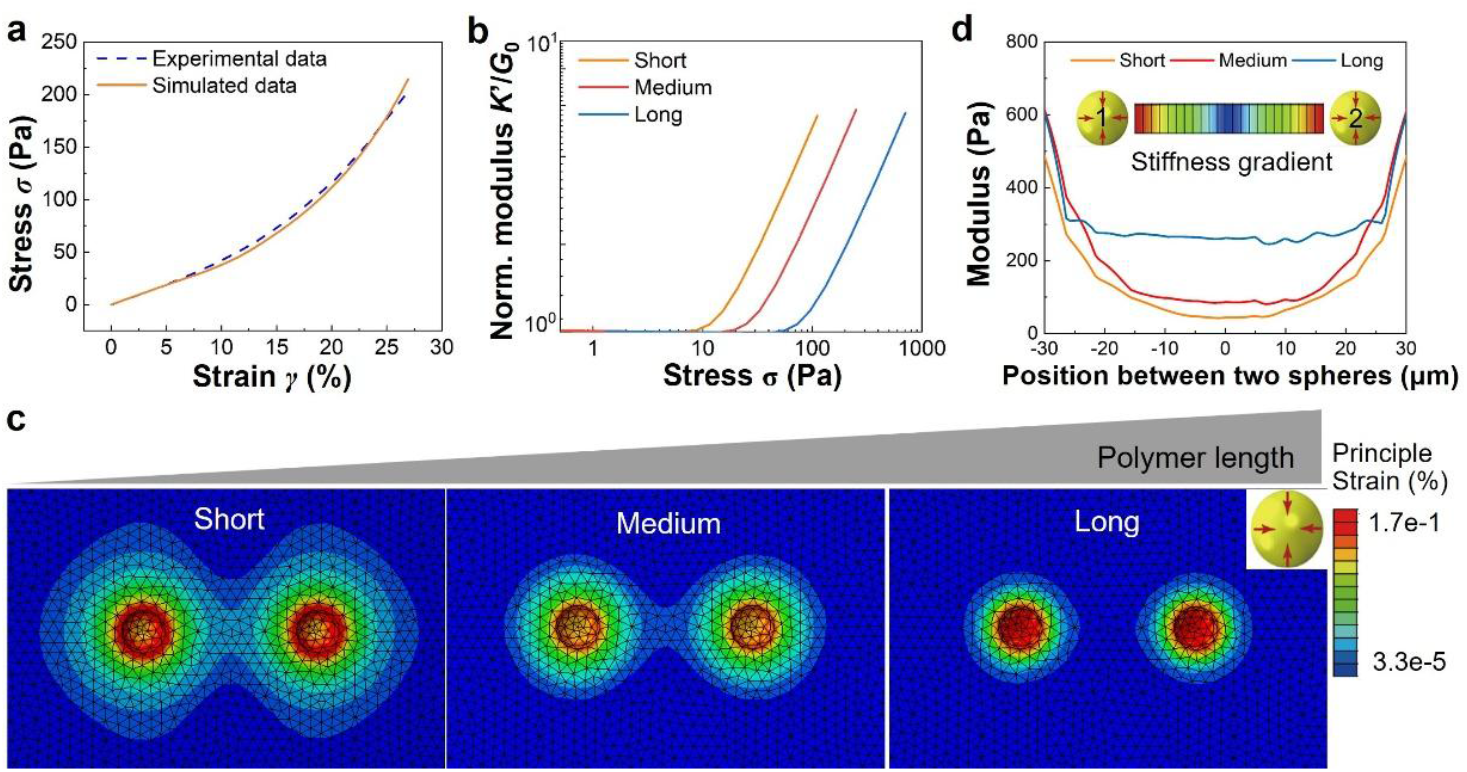
Mechanical modeling of PIC hydrogels reveals the role of matrix stiffening in long-range cell-cell communication. (a) Simulated stress–strain curve of PIC hydrogels closely matches experimental data, validating the constitutive model. Experiments, dashed; Model, solid. (b) Simulated modulus (normalized, *K*′/*G*_0_) as a function of applied stress for short, medium, and long PIC polymers by tuning the critical stretch λ_*c*_, showing an increase in critical stress (*σ*_*c*_) with polymer length. (c) Numerical model of two contractile spherical inclusions embedded in PIC hydrogels with different polymer lengths. Short polymers generate larger high-strain region that overlap, forming aligned high-strain tracts, whereas long polymers restrict strain propagation. (d) Predicted spatial stiffness map along the line connecting sphere 1 and sphere 2 for networks with different polymer lengths. The model shows pronounced pericellular stiffening around each sphere that gradually decays with distance toward the midpoint (0 μm), generating apparent modulus gradients that can mediate long-range mechanical communication.

To probe the effect of polymer length on cell behaviors, we modeled two contractile cells (represented by spherical inclusions) embedded in PIC hydrogels. Remarkably, we found that, in short PIC hydrogels, the pericellular strain fields expanded and overlapped to form a continuous band, indicating that one cell can sense the strain field generated by its neighbor, thereby facilitating long-range cell–cell communication. In contrast, in long PIC hydrogels, the strain fields remained localized with minimal overlap and limited transmission range, leading to negligible communication between cells (**Figure 3**c). Therefore, we conclude that shortening the polymer length (or equivalently reducing critical stretch λ_*c*_) extends the range of force transmission, increases network sensitivity to traction forces, and facilitate the cell spreading and morphological transition, eventually promoting cell-cell long range communication through the matrix.

Furthermore, our model predicted that the apparent modulus 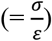 under small strain increased more than 1 order of magnitude in short polymer matrix, i.e., from far-field values (∼68 Pa) to ∼500 Pa adjacent to the cell surface, and decayed gradually with distance (**Figure 3**d). This indicates a substantial cell-induced local stiffening around the cell. Similar trends occur in matrices with medium and long polymers, but the stiffening gradients extend over shorter distances. Note that such stiffening arises from the *W*_*f*_ contribution reflecting a contractility-guided fiber alignment, suggesting the architecture remodeling is more pronounced in short PIC hydrogels. Based on these results, we selected short polymer RGD-functionalized PIC hydrogels for subsequent studies of matrix remodeling and cancer cell invasion.

### Cellular force-induced fiber remodeling and invasion

The natural ECM is defined by its fibrous architecture, and the influence of fiber organization extends beyond primary tumors into metastatic sites. In colorectal cancer, fibrosis of metastatic lymph nodes has been correlated with reduced patient survival^45^, while in ovarian cancer, collagen fiber orientation within metastases is closely associated with disease stage and prognosis^46^. In general, randomly oriented fiber networks are characteristic of normal stroma, whereas aligned and thickened fibers are hallmarks of fibrosis and tumor progression^27,47^. Although engineered hydrogels are widely used to mimic tissue-relevant stiffness and biochemical composition, relatively few recapitulate the in vivo fiber architecture^48-50^. Capturing this architectural dimension is therefore essential for modeling ECM remodeling during cancer invasion and metastasis. PIC hydrogels intrinsically form fibrous networks without additional processing, owing to the semiflexible polymer backbone and hydrophobic assembly that occurs upon warming. PIC polymers can be readily fluorescently labeled, allowing direct visualization of cell–matrix interactions. Imaging revealed that PIC hydrogels form highly heterogeneous fibrous networks with pore sizes of several micrometers, which remain constant when mechanics are tuned by polymer length^51^ (**Figure S5** in the supporting information). During spreading and migration, both MDA-MB-231 cells and CAFs actively remodeled PIC fibers through pulling and pushing, generating pronounced fiber densification and reorganization across the matrix (**Figure 4**a,b). This densification was partially alleviated by treatment with the actin-disrupting agent cytochalasin D, confirming that cell-generated tension drives the process (**Figure S6** in the supporting information). The inherent plasticity of PIC ensures that such cellular forces lead to permanent matrix deformations (**Figure 1**c). Interestingly, we found that cells were able to invade through PIC hydrogels in a Transwell format, in which PIC/cell constructs were directly placed on the insert instead of conventional Matrigel, taking advantage of PIC’s thermoresponsive gelation. After 3 days, large numbers of MDA-MB-231 cells migrated through the PIC network, while CAFs invaded even more rapidly, within 1 day, consistent with their strong contractile capacity (**Figure 4**c-f). Importantly, RGD functionalization was essential for cell–matrix engagement. Because PIC is not enzymatically degradable, the observed fiber remodeling and invasion must arise from purely mechanical mechanisms, consistent with an MMP-independent mode of migration.

**Figure 4.**
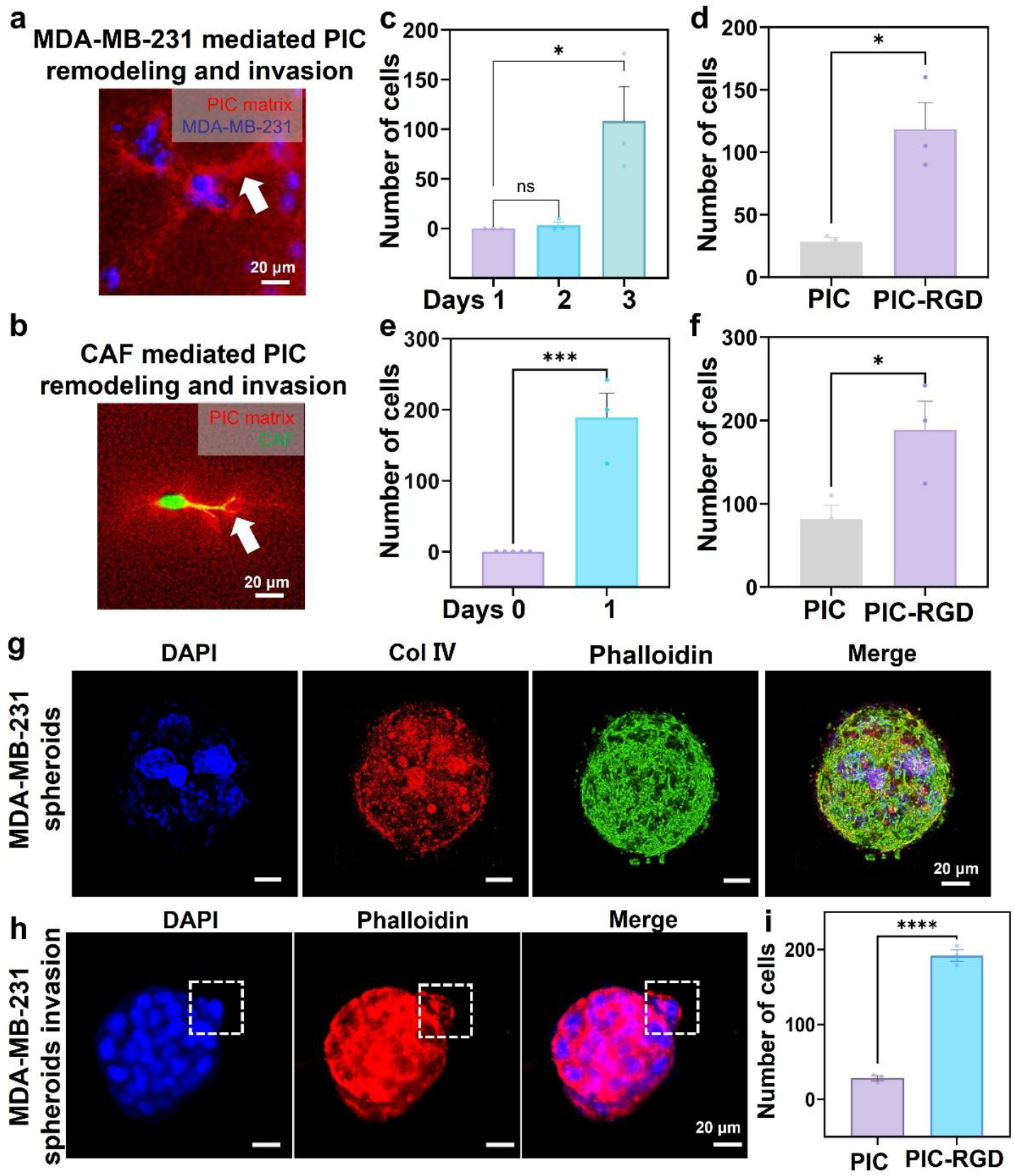
PIC hydrogels support cellular force-mediated fiber remodeling and invasion independent of enzymatic degradation. Representative fluorescence image of PIC fiber rearrangement mediated by MDA-MB-231 cells on PIC hydrogels (a) and CAFs in PIC hydrogels (b). White arrows indicate regions of fiber densification with increased fluorescence intensity. Red: PIC fibers. Blue: MDA-MB-231 cells (CellTracker). Green: CAF-GFP. (c) Quantification of invading MDA-MB-231 cells in PIC hydrogel-based Transwell assay over 3 days shows progressive invasion. (d) RGD functionalization significantly increases the number of invading MDA-MB-231 cells compared with non-functionalized PIC. (e) Quantification of CAF invasion shows rapid penetration within 1 day. (f) CAF invasion is enhanced in RGD-functionalized PIC compared with non-functionalized PIC. (g) Immunostaining of MDA-MB-231 spheroids reveals a compact morphology enclosed by a collagen IV–rich basement membrane (red), with DAPI (blue) and phalloidin (green). (h) Confocal images show invasion of MDA-MB-231 spheroids through the PIC matrix, with actin-rich protrusions emerging from the spheroid periphery. (i) Quantification confirms that invasion from spheroids is significantly higher in RGD-functionalized PIC compared with non-functionalized PIC. All experiments were performed in short PIC hydrogels at 1.0 mg mL ^−1^, corresponding to a low critical stress and high plasticity. Data represent mean ± SEM (n = 3–5 biological replicates). ns, not significant; \**p* < 0.05, \*\*\**p* < 0.001, by One-way ANOVA Tukey’s test. Scale bars = 20 μm.

To model collective migration within the tumor microenvironment, we next examined the invasion of MDA-MB-231 cells under spheroid conditions. After 7 days of culture, MDA-MB-231 cells formed compact spheroids enclosed by a basement membrane, with collagen IV serving as a barrier that maintains spheroid integrity (**Figure 4**g). These spheroids were subsequently encapsulated and subjected to the Transwell assay. As shown in **Figure 4**i, MDA-MB-231 cells breached the basement membrane and invaded through the surrounding matrix within 5 days, a process that occurred approximately two days later than single-cell invasion (**Figure 3**c). Notably, sprouting structures were observed at the spheroid periphery (**Figure 4**h), directly demonstrating that groups of cells can coordinate their migration and invade collectively.

### CAFs promote cancer metastasis through physical guidance

Both biochemical signaling and mechanical contractility of CAFs are recognized as key drivers of cancer cell invasion and metastasis. Yet, in vitro models that faithfully capture the complexity of CAF–matrix–cancer cell interactions remain limited in synthetic systems^27^. To address this, we established a co-culture system in PIC hydrogels to investigate how CAFs influence cancer cells through fiber remodeling. As shown in **Figure 5**a, MDA-MB-231 cells and CAFs were encapsulated together within PIC constructs. CAFs rapidly spread and adopted elongated morphologies consistent with their strong contractile forces, whereas MDA-MB-231 cells predominantly formed spheroids after 7 days. Strikingly, in the presence of CAFs, MDA-MB-231 spheroids lost compactness and cohesion, developing irregular surfaces with diffuse, outward-spreading fronts (**Figure 5**b). Fiber staining revealed that CAFs remodeled the surrounding PIC network and established aligned, densified tracts oriented toward the spheroids (**Figure 5**c). These physical tracks provided directional cues, enabling CAFs to mechanically communicate with cancer cells at a distance and to promote invasive behavior.

**Figure 5.**
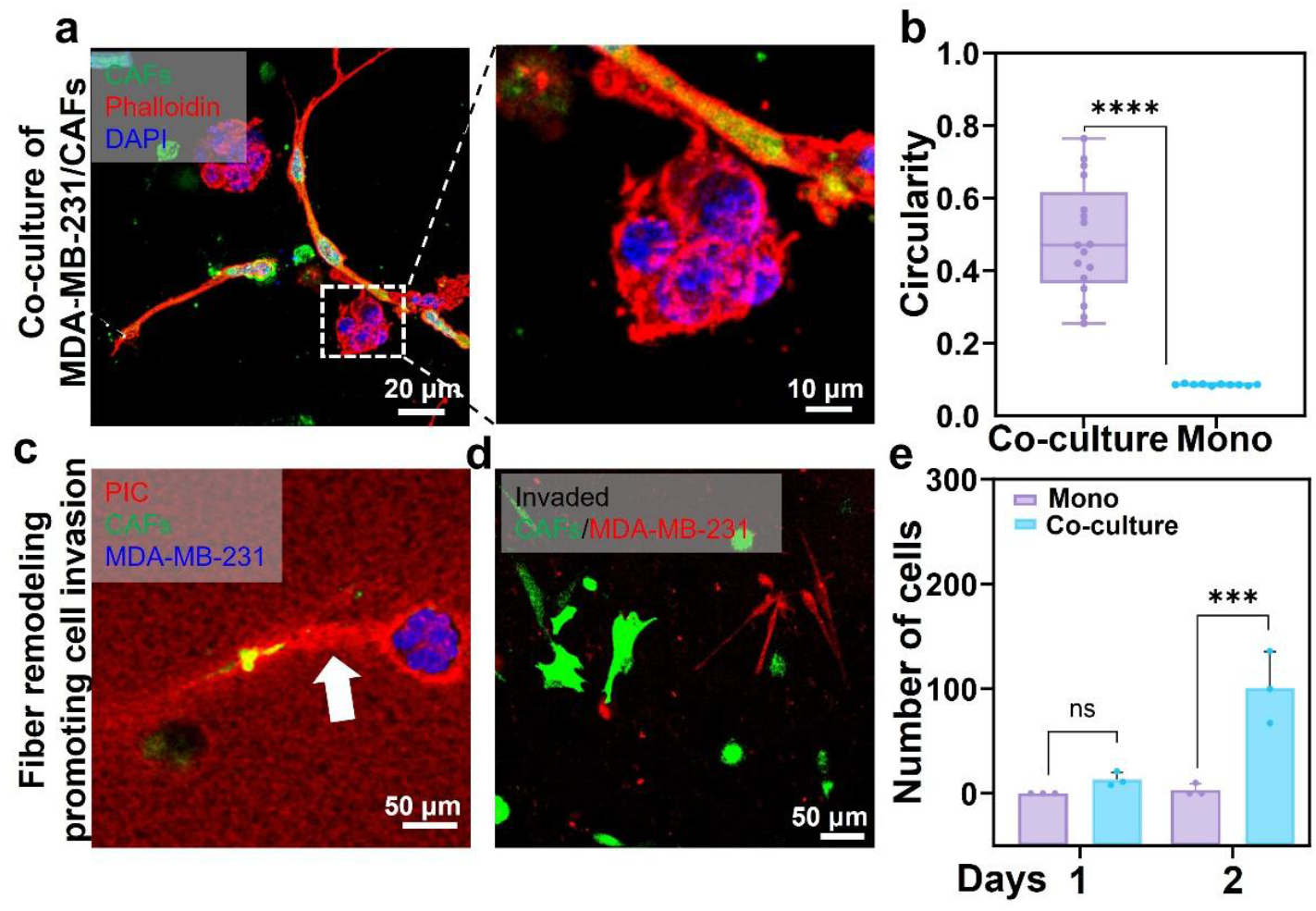
CAFs promote cancer cell invasion in PIC hydrogels through physical fiber remodeling. (a) Confocal images of co-cultures of MDA-MB-231 cells (nuclei, DAPI, blue; F-actin, phalloidin, red) with GFP-expressing CAFs (green) in PIC hydrogels after 7 days. Enlarged image (right) shows direct CAF–cancer cell contact. (b) Quantification of circularity demonstrates that MDA-MB-231 cells in co-culture adopt more irregular and invasive morphologies compared with monoculture. n = 18-20. (c) Representative image showing CAF-mediated fiber remodeling (white arrow) guiding MDA-MB-231 cell spheroids within the PIC matrix. Red: PIC fibers. Blue: MDA-MB-231 cells (CellTracker). Green: CAF-GFP. Fluorescence image (d) and quantification (e) of MDA-MB-231 cells (CellTracker, red) and CAFs (GFP, green) invaded through PIC hydrogels into the bottom chamber in Transwell assays, showing significantly accelerated invasion in co-culture compared with monoculture after 2 days. All experiments were performed in short PIC hydrogels with GRGDS functionalization at 1.0 mg mL ^−1^. Data represent mean ± SEM (n = 3–5 biological replicates) ns, not significant; \*\*\**p* < 0.001, \*\*\*\**p* < 0.0001, by One-way ANOVA Tukey’s test. Scale bars: 20 μm (a), 10 μm (a, zoom), 50 μm (c, d).

The matrix structural alterations stiffen the pericellular environment and enrich adhesive ligands, thereby promoting cancer cell proliferation and strengthening cell– matrix interactions. In alignment with our findings, histological analyses of human breast carcinoma have documented collectively invading cells that migrate along preformed microtracks within aligned collagen fibers^52^. Previous studies have further shown that both collective invasion and collagen alignment are associated with poor prognosis and increased metastatic potential in breast cancer^53,54^. In squamous cell carcinoma, fibroblast-mediated remodeling also facilitates invasion, as stromal fibroblasts deposit fibronectin and tenascin-C to generate aligned tracks that guide directional cancer cell migration^55^. To our knowledge, this represents the first in vitro demonstration of CAF-mediated physical fiber remodeling driving cancer invasion in a fully synthetic matrix. We further adapted this co-culture system to a Transwell assay. As expected, MDA-MB-231 cells invaded more rapidly in the presence of CAFs than in monoculture. While cancer cells alone required more than 3 days to breach the PIC matrix, co-cultures achieved invasion within only 2 days (**Figure 5**d,e).

### Applying cell-matrix mechanical interaction models for drug evaluation

Marimastat and Tanomastat are broad-spectrum MMP inhibitors that once showed considerable promise as anticancer agents but ultimately failed in clinical trials^11,13^. Notably, both compounds often appear effective in conventional invasion assays using Matrigel or collagen, suggesting that traditional platforms may overestimate their therapeutic potential. Given the persistently high attrition rates in oncology drug development, there is a pressing need for advanced 3D systems that more accurately predict patient outcomes^56,57^. Such models must recapitulate disease-relevant mechanisms, including tissue-specific phenotypes and the dual biochemical and mechanical pathways of cell–matrix interaction. By capturing both dimensions, they may reveal targetable processes that are overlooked in standard assays and improve the translational success of preclinical findings.

It is well established that cancer cells can invade through two principal strategies, i.e., protease-mediated degradation and force-driven matrix remodeling. Yet, most drug evaluation platforms primarily emphasize biochemical pathways, leaving the role of biophysical cues largely untested. To address this gap, we extended our PIC-based cell–matrix interaction model to drug evaluation. In this assay, a thin layer of PIC–cell constructs were placed on Transwell inserts with a chemoattractant in the lower chamber, and candidate drugs were added to the upper chamber (**Figure 6**b). Strikingly, neither Marimastat nor Tanomastat suppressed invasion in PIC-based models, despite their efficacy in Matrigel (**Figure 6**c). This simple but direct experiment underscores the importance of including physical remodeling in drug evaluation. While it remains true that tumor cells secrete diverse proteases and that therapeutic efficacy may depend on optimized treatment timing^10^, these results demonstrate that mechanical models can complement biochemical assays, providing a more comprehensive and predictive framework for preclinical drug testing.

**Figure 6.**
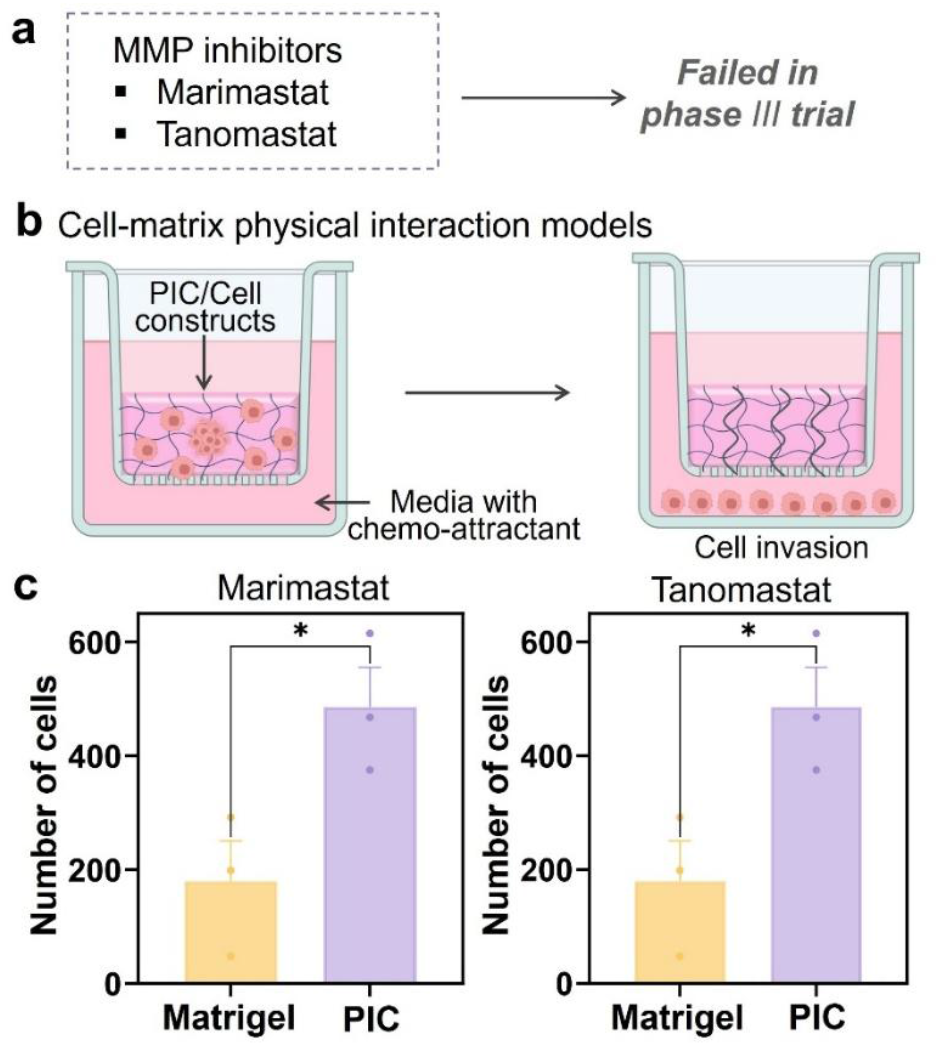
PIC-based cell-matrix mechanical interactions models for drug evaluation. (a) Broad-spectrum MMP inhibitors Marimastat and Tanomastat showed promise in preclinical studies but failed in phase III clinical trials. (b) Schematic of the PIC-based Transwell invasion assay. PIC/cell constructs were placed on inserts with chemoattractant in the lower chamber. In this system, invasion occurs through force-mediated fiber remodeling rather than enzymatic degradation. (c) Quantification of invading MDA-MB-231 cells in Matrigel-versus PIC-based Transwell assays in the presence of Marimastat (left) or Tanomastat (right). Data are shown as mean ± SEM (n = 3 biological replicates, \**p* < 0.05, by one-way ANOVA with Tukey’s test).

## Discussion

Most current invasion and drug evaluation platforms emphasize biochemical pathways while overlooking the physical strategies that cells use to navigate 3D tissues^56^. In fibrous nonlinear matrices, traction forces can be transmitted, focused, and amplified across many cell diameters through tension-driven fiber recruitment and alignment^27^. This phenomenon, captured in both continuum and network models, has been widely observed in collagenous tissues. PIC hydrogels reproduce these mechanics and, uniquely, allow control of the nonlinear regime and force sensitivity through polymer length. As such, PIC offers a rare combination of physiological relevance for force-guided migration with rigorous control over matrix parameters. Moreover, its thermoreversible gelation enables straightforward adaptation to Transwell assays and direct visualization of fiber architecture, invasion fronts, and drug responses, without the compositional variability that complicates tumor-derived matrices.

Mechanistically, our findings demonstrate that CAFs enhance invasion by physically remodeling the matrix. CAFs densify and align fibers, generate directional tracks that guide collective egress from spheroids, and accelerate invasion in the Transwell assay. These changes increase the apparent pericellular modulus and create stiffness gradients that extend between neighboring cells. Such behavior aligns with theoretical predictions of long-range force transmission and multiaxial coupling in fibrous networks. Our continuum mechanics framework, parameterized by PIC’s plateau modulus *G*_0_ and critical stress *σ*_c_, provides a mechanistic explanation. A lower recruitment threshold 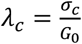 enlarges stiffening halos and strengthens intercellular coupling. This suggests that in tissue, protease-mediated degradation and force-guided fiber remodeling may act synergistically to promote invasion.

From a translational perspective, our work highlights the limitations of traditional assays for drug evaluation. Broad-spectrum MMP inhibitors that block invasion in degradable matrices failed in PIC, where the absence of enzymatic cleavage sites forces invasion to proceed through force-mediated plastic remodeling. This finding provides a plausible explanation for the weak clinical performance of these agents, despite encouraging preclinical data in Matrigel and collagen systems. More broadly, it argues for the development of assay platforms that integrate both biochemical and biophysical routes of invasion, with the goal of improving predictive power in the evaluation of antimetastatic therapies.

## Materials and Methods

### Materials

PIC polymers with defined contour lengths were synthesized by varying the total monomer-to-catalyst ratios of 2,000:1 and 5,000:1, corresponding to short and long polymers, respectively, following established protocols^39^. For biofunctionalization with cell-adhesive peptides, the polymerization feed consisted of 1 mol% azide-terminated monomers and 99 mol% methoxy-terminated monomers. Unless otherwise specified, PIC hydrogels were prepared in sterile phosphate-buffered saline (PBS, pH 7.4). DBCO-PEG_4_-GRGDS peptides were custom-synthesized by Shanghai ABR Pharm Co., Ltd. CellTracker™ Deep Red, phalloidin, and DAPI were purchased from Thermo Fisher Scientific. DBCO-PEG_4_-TAMRA, Cytochalasin D, Marimastat, and Tanomastat were obtained from Sigma-Aldrich. Anti-Collagen IV antibody (ab6586) was obtained from Abcam. μ-Slide 15 Well 3D plates were from ibidi.

### Biofunctionalization of PIC polymers

A total of 35 mg of azide-functionalized PIC was completely dissolved in 14 mL of acetonitrile and stirred for 6 h at room temperature. Afterward, a DBCO-peptide solution was prepared by dissolving 1 mg of DBCO-PEG_4_-GRGDS peptide in 83 μL of DMSO. The peptide solution was then added dropwise to the PIC solution under gentle stirring, and the reaction was allowed to proceed overnight at room temperature. The reaction mixture was precipitated by adding isopropyl ether at a 1:10 (v/v) ratio and centrifuged to collect the biofunctionalized PIC-GRGDS polymer and air-dried for 2 days. Same procedure for PIC-HVADI functionalization.

### Fluorescence labeling of PIC hydrogels

To fluorescently label the PIC polymers, a red-emitting dye, DBCO-PEG_4_-TAMRA, was conjugated to the azide groups on the polymer via strain-promoted azide–alkyne cycloaddition. The dye was added to the PIC solution at a ratio of 1 mg polymer : 0.5 nmol dye, incubated on ice for 5 min, and subsequently used for cell encapsulation.

### Rheological Characterization

The mechanical properties of PIC hydrogels were measured using a stress-controlled rheometer (Discovery HR-2, TA Instruments) equipped with an aluminum parallel-plate geometry. Samples were loaded as cold polymer solutions (kept on ice), and the temperature was increased from 5 °C to 37 °C at a rate of 1.0 °C min^-1^.

#### Storage modulus (G′)

Oscillatory shear tests were performed at a strain amplitude of *γ* = 0.04 and frequency of *ω* = 1.0 Hz. The gelation temperature was defined as the point at which *G*′ began to increase with rising temperature.

#### Strain-stiffening behavior

Nonlinear mechanical response was evaluated using a prestress protocol^34^. A constant prestress (*σ*_0_) was applied while superimposing a small oscillatory stress (*σ* < 0.1 *σ*_0_) at frequencies between 10 and 0.1 Hz at 37 °C. The differential modulus (*K*′), defined as *K*′ = *∂σ/∂γ*, was plotted as a function of applied prestress at *ω* = 1.0 Hz to determine the critical stress (*σ*_c_) marking the onset of strain stiffening.

#### Plasticity measurement

Creep and recovery tests were performed using a rheometer equipped with a 25 mm parallel-plate geometry and a 500 μm gap. The polymer solution was pre-cooled and then loaded onto the rheometer stage. After equilibration at a frequency of *ω* = 1.0 Hz and a strain amplitude of *γ* = 0.04, ensuring a stable storage modulus, the creep–recovery test was initiated. During the creep phase, a constant stress of 10 Pa was applied for 300 s, and the resulting strain evolution of the hydrogel was continuously recorded to monitor deformation under sustained load. In the recovery phase, the applied stress was removed, and the residual strain was tracked until a new equilibrium was reached. The irreversible component of the strain provided a quantitative measure of the plasticity of the hydrogel.

### Cell encapsulation

Dry PIC polymers were sterilized under UV light for 20 min, then dissolved in cell culture medium at 4 °C for 24 h to reach the desired concentration, ensuring complete dissolution. The cell suspension and PIC solution were mixed at a 1:1 (v/v) ratio on ice and transferred into μ-Slide plates. The constructs were incubated at 37 °C to induce gelation and form stable cell/gel constructs. The final cell density was 1 × 10^5^ cells/mL. All media were pre-warmed to 37 °C before use, and the culture medium was refreshed every 3 days.

### Co-culture of CAFs and MDA-MB-231 Cells

MDA-MB-231 and CAF-GFP cells were harvested as single-cell suspensions and mixed at a 1:2 ratio (MDA-MB-231 : CAF-GFP). The mixed cell suspension was combined 1:1 (v/v) with PIC hydrogel (1 mg/mL), and 10 μL of the resulting mixture was seeded into μ-Slide plate. The constructs were incubated at 37 °C for 20 min to allow complete gelation and then maintained in complete culture medium, which was refreshed every 3 days. After 7 days of culture, samples were fixed and stained for confocal imaging.

### PIC-based Transwell assay

#### Single-cell invasion assay

Cells were resuspended at a density of 2 × 10^5^ cells mL^-1^ and mixed 1:1 (v/v) with short PIC hydrogel (1 mg mL^-1^). A total of 75 μL of the cell/hydrogel mixture was added to the upper chamber of Transwell inserts (8 μm pore size, 24-well format). Inserts were placed into 24-well plates and incubated at 37 °C, 5% CO_2_ for 30 min to allow complete gelation. Serum-free medium was then added to the upper chamber, while complete medium containing 10% FBS was added to the lower chamber to establish a chemoattractant gradient. After 72 h, cells were fixed with 4% paraformaldehyde (15 min) and stained with 0.1% crystal violet (30 min). Invaded cells on the underside of the membrane were imaged by bright-field microscopy, and invasion was quantified using ImageJ.

#### Spheroid invasion assay

To generate spheroids, MDA-MB-231 cells were encapsulated in PIC hydrogels functionalized with GRGDS and cultured for 7 days to form compact aggregates. Spheroids were recovered by cooling the plate on ice to liquefy the thermoresponsive gel and gently collected by centrifugation. The recovered spheroids were remixed 1:1 (v/v) with fresh PIC hydrogel and seeded into Transwell inserts (8 μm pores). After gelation at 37 °C for 30 min, serum-free medium was added to the upper chamber and complete medium with 10% FBS to the lower chamber. After 5 days, invaded spheroids and migrated cells were fixed in 4% paraformaldehyde and stained with phalloidin to visualize actin structures.

#### CAF–GFP and MDA-MB-231 co-culture invasion assay

MDA-MB-231 cells were pre-labeled with CellTracker™ Deep Red for 30 min and then mixed with CAF-GFP cells at a 2:1 ratio (CAF-GFP : MDA-MB-231) to obtain a homogeneous suspension. A total of 75 μL of the mixture was added to Transwell inserts (8 μm pores) and incubated at 37 °C for 30 min to induce gelation. Serum-free medium was supplied to the upper chamber and complete medium containing 10% FBS to the lower chamber. After 72 h, invaded cells were imaged by confocal microscopy, recording both the red (MDA-MB-231) and green (CAF-GFP) fluorescence channels.

### Imaging of cell-induced PIC fiber remodeling

Fluorescently labeled PIC-TAMRA polymers were used to encapsulate MDA-MB-231 (CellTracker™ Deep Red) and CAF-GFP cells for visualization of fiber remodeling. After 1 day of encapsulation, constructs were imaged using confocal microscopy to assess fiber organization around cells.

To examine the contribution of cellular contractility, cytochalasin D (5 μM) was added to disrupt actin filaments and reduce intracellular tension. After 20 min of treatment, the same fields of view were re-imaged to compare fiber morphology before and after cytoskeletal relaxation.

### Immunofluorescence analysis

For immunofluorescence staining, cells were encapsulated in PIC hydrogels and cultured in μ-Slide chambers for 7 days. All staining solutions were pre-warmed to 37 °C prior to use. Samples were washed twice with PBS and fixed in 4% paraformaldehyde (PFA) for 1 h at 37 °C. After fixation, residual formaldehyde was quenched with 0.1 M glycine in PBS for 5 min, followed by two additional PBS washes.

Samples were then permeabilized with 0.1% Triton X-100 at 37 °C for 30 min and blocked with 1% goat serum overnight at 37 °C. Without intermediate washing, constructs were incubated with anti–collagen IV primary antibody at 37 °C overnight, followed by two washes in PBS (1 h each). Subsequently, goat anti-rabbit secondary antibody was added and incubated for 4 h at 37 °C.

For actin cytoskeleton staining, samples were treated with phalloidin (5 μg mL^-1^) for 90 min at 37 °C, followed by DAPI staining for 15 min. Samples were washed three times with PBS (5 min each) before imaging. Fluorescence images were acquired using a confocal laser scanning microscope (Olympus FV1000, Japan).

### Statistical Analysis

Statistical analyses were performed using one-way ANOVA followed by Tukey’s multiple-comparison test in SPSS. Data are presented as mean ± SEM. Statistical significance was defined as * *P* < 0.05, ** *P* < 0.01, *** *P* < 0.001.

### Numerical simulations

We implemented the PIC constitutive model in ABAQUS via a user-defined material (UMAT) subroutine. The domain was discretized with C3D10 (quadratic tetrahedral) elements to investigate the effects of material nonlinearity (strain stiffening). The continuum-based model employs the specified free-energy density to capture nonlinear strain-stiffening behavior. The derivations of tangent modulus tensor in the material description C^SC^, the tangent modulus tensor for the convected rate of the Kirchhoff stress C^τC^, the tangent modulus tensor for the Jaumann rate of the Kirchhoff stress C^τj^, and the material Jacobin C^MJ^ (needed for the user material model) can be found in Ref^32^.

#### Shear model of PIC

A schematic of the simple-shear model is shown in Figure S7. A regular hexahedron with a side length of 100 *µm* was fully fixed at the bottom surface, and the top surface was fixed in x and z but a horizontal rightward displacement u was applied in y. Then we extracted the Cauchy shear stress and shear strain in the deformed model, and obtained the stress-strain curve of the constitutive model with a user-defined post-processing program.

#### Modeling of cell-cell long-range communications

he long-range communication model between cells describes the long-range mechanical signal transmission under the action of cell contractile force in PIC. This model is a rectangular prism with a length of 200 µm, a width of 100 µm, and a height of 50 µm respectively. For the bottom surface, the nodes were fixed in x, y, and z. For the top surfaces, there were two hemispherical cavities with a diameter of 20 µm (representing cells), which were 100 µm apart. A total force of 31.4 nN was added to the inner hemisphere, which was equivalent to a cell traction stress of about 100 Pa. The maximum principal stress and principal strain data of all nodes between the two hemispheres on the upper surface of the deformed model were first extracted and analyzed with a post-processing program to obtain the spatial map of modulus change.

## Acknowledgments

This project has received funding from the National Natural Science Foundation of China (32471368, 22207029, 22077025, and 12402195), Natural Science Foundation of Hebei Province (No. B2025202003), Financial Support Project of Central Government for Promoting Development of Science and Technology of Hebei Province (No. 226Z2401G), the Program for Overseas Researchers of Hebei Province (C20230503), the Fonds Wetenschappelijk Onderzoek (12A2423N), Shenzhen Science and Technology Innovation Commission (GXWD2023113014055503), Guangdong Basic and Applied Basic Research Foundation (2023A1515110909), and Department of Science and Technology of Guangdong Province (2023QN10C056). Figure 1 was created with support of BioRender.

