## Supporting Information for "3D Cell-Matrix Mechanical Interaction Models for Cancer Invasion and Drug Evaluation"

**Constitutive model of PIC hydrogel**

The total Helmholtz free-energy density is expressed as[1]

$$W = W_b + W_f$$

with the first term being Neo-Hookean isotropic response

$$W_b = \frac{\mu_0}{2}\left(\overline{I_1} - 3\right) + \frac{\kappa}{2}(J-1)^2$$

where $\mu_0$ and $\kappa$ are the shear and bulk modulus, respectively. $J = \det(\mathbf{F})$ with $\mathbf{F}$ being the deformation gradient, and $\overline{I_1} = tr(\overline{\mathbf{C}})$ is the first invariant of the isochoric part of right Cauchy–Green tensor $\overline{\mathbf{C}} = \mathrm{J}^{-2/3}\mathbf{C}$, with $\mathbf{C} = \mathbf{F}^{\mathrm{T}}\mathbf{F}$. The fibrous contribution $W_f$ sums along three principal stretch directions $\lambda_i$ with a tension-only response:

$$W_f = \sum_{i=1}^{3} f(\lambda_i)$$

$$\frac{\partial f(\lambda_i)}{\partial \lambda_i} = \begin{cases} 0, & \lambda_i \le \lambda_1 \\ \dfrac{E_f \left(\dfrac{\lambda_i - \lambda_1}{\lambda_2 - \lambda_1}\right)^n (\lambda_i - \lambda_1)}{n+1} & \lambda_1 < \lambda_i < \lambda_2 \\ E_f\left[\dfrac{\lambda_2 - \lambda_1}{n+1} + \dfrac{(1 + \lambda_i - \lambda_2)^{m+1} - 1}{m+1}\right], & \lambda_i > \lambda_2 \end{cases}$$

This recruitment law enforces (i) no fibrous stress below a critical stretch $\lambda_c$ (isotropic response) and (ii) strain-stiffening above $\lambda_c$ with small-strain fiber modulus $E_f$ and stiffening index $m$. We ensure smooth tangents with a narrow transition band of width $\lambda_t = 0.25\lambda_c$, using $\lambda_1 = \lambda_c - \lambda_t/2$, $\lambda_2 = \lambda_c + \lambda_t/2$, and $n = 5$.

For semiflexible PIC bundle networks, the plateau modulus $G_0$ and critical stress $\sigma_c$

relate to molecular geometry as[2]

$$G_0 = 6\frac{l_M N_A}{M}\frac{c}{N}k_B T\frac{l_p^2}{l_c^3}$$

$$\sigma_c = \frac{l_M N_A}{M}\frac{c}{N}k_B T\frac{l_p}{l_c^2}$$

where $c$ is polymer concentration, $N$ chains per bundle, $l_{p,B}$ bundle persistence length, and $l_c$ polymer contour length, eliminating constants, yields a link between the recruitment threshold and molecular geometry:

$$\lambda_c = \frac{\sigma_c}{G_0} = \frac{l_c}{6l_p}$$

We set $E_f = G_0$ at small strain and fitted $m$ to capture the nonlinear slope (here, $m$ = 15). The critical stretch $\lambda_c$ was calculated as $\lambda_c = l_c/6l_p$, where $l_p$ is the polymer persistence length, with a mean value of 460 nm[2]. Consequently, variations in the polymer contour length $l_c$ (77 nm, 105 nm, and 218 nm, corresponding to short, medium, and long polymers, respectively, as reported in our previous work[3]), shift both $\sigma_c$ and $\lambda_c$, and thereby modulating the onset of fiber recruitment without altering the overall network topology.

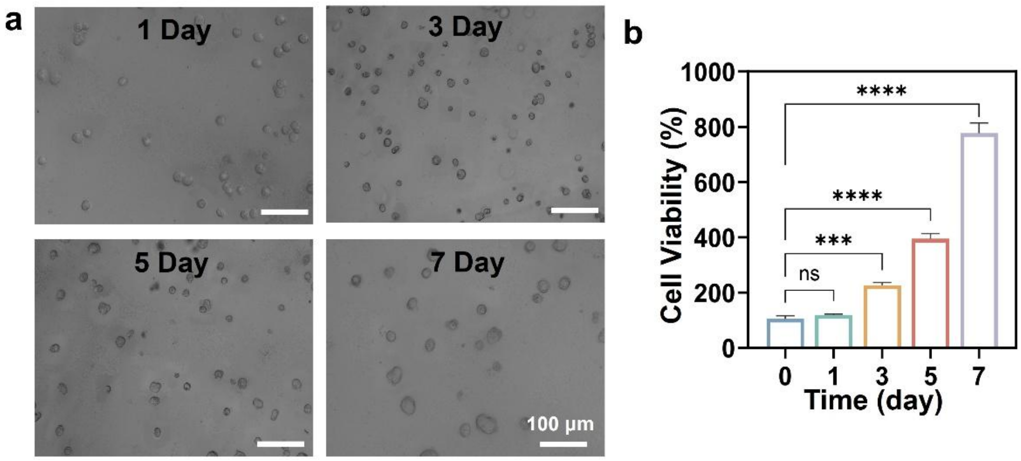


**Figure S1**. **Cell proliferation and viability of MDA-MB-231 cells encapsulated in PIC hydrogels over time.** (a) Representative bright-field images showing MDA-MB-231 cells cultured in PIC hydrogels for 1, 3, 5, and 7 days, illustrating gradual cell proliferation and aggregation into small clusters. Scale bar = 100 μm. (b) Quantitative analysis of cell viability (%) at different time points, showing a significant increase in viability from day 3 onward, indicating good cytocompatibility and sustained cell growth within PIC hydrogels. PIC hydrogels were functionalized with GRGDS at a polymer concentration of 1.0 mg $mL^{-1}$, corresponding to a GRGDS density of 31.4 μM. Data are presented as mean ± SEM (n = 3); ns, not significant; ***$p < 0.001$; ****$p < 0.0001$ (one-way ANOVA with Tukey's post hoc test).

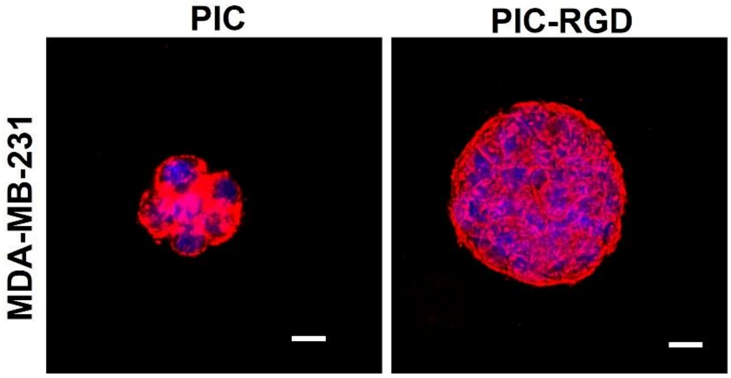


**Figure S2**. **Effect of biofunctionalization on MDA-MB-231 spheroid formation in PIC hydrogels.** Representative confocal fluorescence images of MDA-MB-231 cells cultured for 7 days in non-functionalized PIC, and GRGDS-functionalized PIC (PIC-RGD) hydrogels. Short PIC polymers were used with a concentration of 1.0 mg $mL^{-1}$. F-actin (phalloidin, red); nuclei (DAPI, blue). Scale bar = 20 µm.

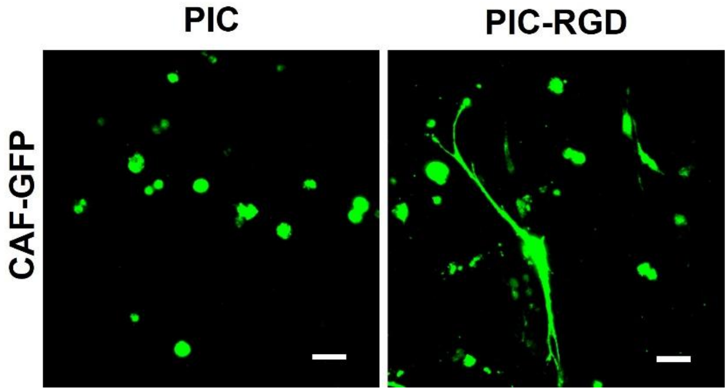


**Figure S3**. **CAF morphology in PIC hydrogels with different biofunctionalizations.** Representative confocal fluorescence images of CAF-GFP cells cultured for 1 day in non-functionalized PIC, and GRGDS-functionalized PIC (PIC-RGD) hydrogels. Short PIC polymers were used with a concentration of 1.0 mg $mL^{-1}$. GFP fluorescence marks CAFs (green). Scale bar = 20 µm.

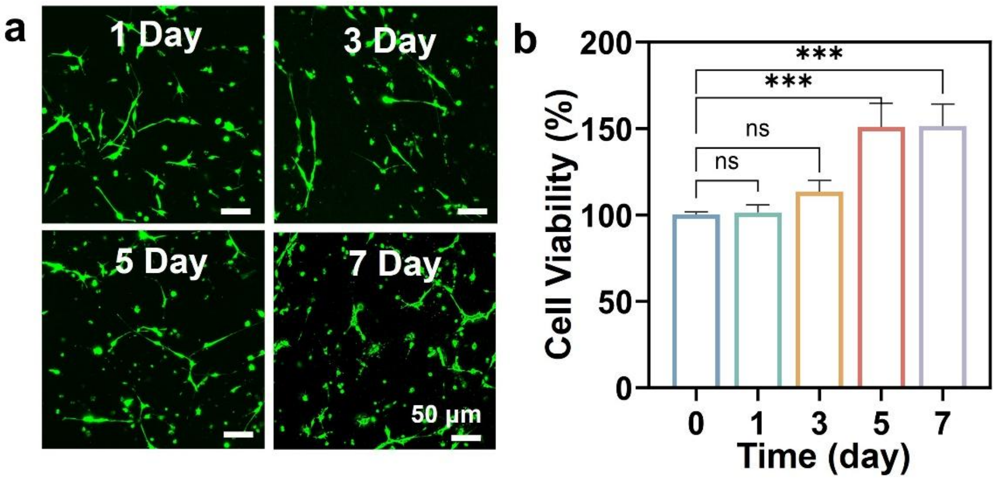


**Figure S4**. **CAFs' viability in PIC-RGD hydrogels as a function of time.** (a) Representative confocal fluorescence images of GFP-expressing CAFs (green) cultured in short PIC-RGD hydrogels for 1, 3, 5, and 7 days, showing progressive cell spreading and network formation over time. Scale bar = 50 μm. (b) Quantification of CAF viability (%) at different time points indicates sustained cell survival and significant proliferation from day 3 onward, confirming the cytocompatibility of the PIC-RGD hydrogel. Short PIC polymers were used with a concentration of 1.0 mg $mL^{-1}$. Data are presented as mean ± SEM (n = 3); ns, not significant; ***$p < 0.001$ (one-way ANOVA with Tukey's post hoc test).

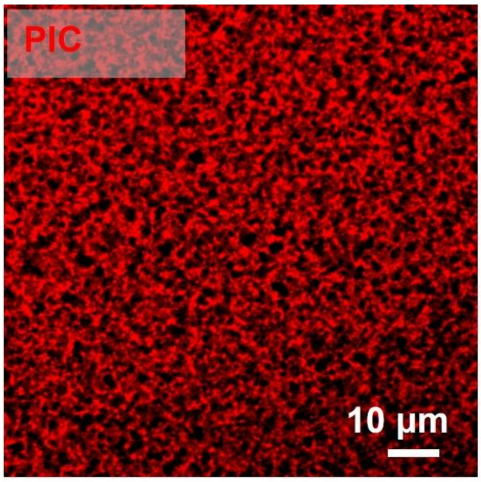


**Figure S5**. **Confocal fluorescence image of the fibrous architecture of PIC hydrogel.** Representative confocal image of fluorescently labeled PIC hydrogel (TAMRA, red) showing a highly porous and heterogeneous fibrous network with micron-scale pore sizes. Short PIC polymers were used with a concentration of 1.0 mg $mL^{-1}$. Scale bar = 10 μm.

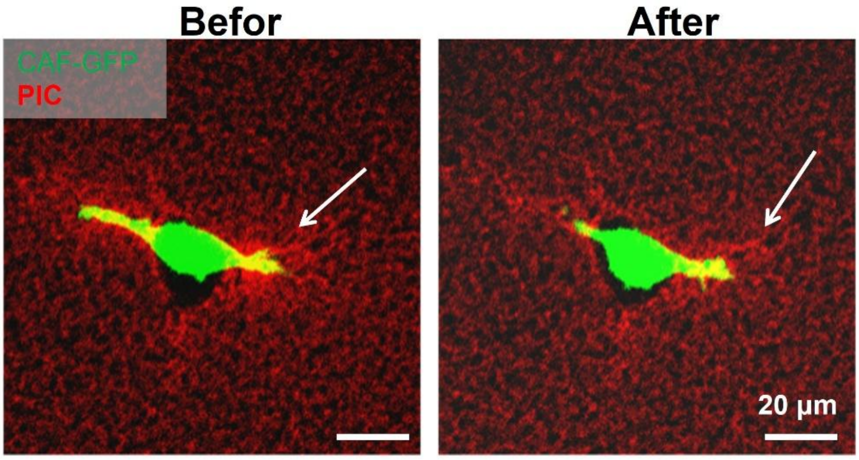


**Figure S6**. **CAF-mediated PIC fiber remodeling before and after releasing cellular forces.** Representative confocal fluorescence images showing CAF-GFP cells (green) embedded in fluorescently labeled PIC hydrogels (red) before and after treatment with 5 μM cytochalasin D. Prior to treatment, CAFs induce local fiber densification and alignment near the cell edge (white arrows). After actin disruption, the densified fiber structure partially relaxes, indicating that CAF-generated contractile forces drive fiber remodeling in the PIC matrix. Short PIC polymers were used with a concentration of 1.0 mg mL$^{-1}$. Scale bar = 20 μm.

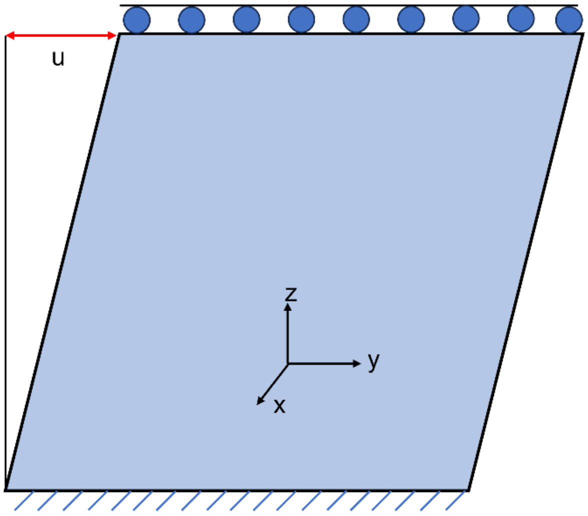


**Figure S7. Simple-shear model for the bulk response of PIC.** A 100-μm cube is modeled; the bottom face is fully fixed, the top face is fixed in the x and z directions and subjected to a uniform rightward displacement in y, lateral faces are traction-free. The macroscopic shear stress is obtained from the reaction force on the top face.
